# A Feedback Mechanism Regulates *Odorant Receptor* Expression in the Malaria Mosquito, *Anopheles gambiae*

**DOI:** 10.1101/2020.07.23.218586

**Authors:** Sarah E. Maguire, Ali Afify, Loyal A. Goff, Christopher J. Potter

## Abstract

Mosquitoes locate and approach humans (‘host-seek’) when specific Olfactory Neurons (ORNs) in the olfactory periphery activate a specific combination of glomeruli in the mosquito Antennal Lobe (AL). We hypothesize that dysregulating proper glomerular activation in the presence of human odor will prevent host-seeking behavior. In experiments aimed at ectopically activating most ORNs in the presence of human odor, we made a surprising finding: ectopic expression of an *AgOr (AgOr2)* in *Anopheles gambiae* ORNs dampens the activity of the expressing neuron. This contrasts studies in *Drosophila melanogaster*, the typical insect model of olfaction, in which ectopic expression of non-native ORs in ORNs confers ectopic neuronal responses without interfering with native olfactory physiology. To gain insight into this dysfunction in mosquitoes, RNA-seq analyses were performed comparing wild-type antennae to those ectopically expressing *AgOr2* in ORNs. Remarkably, almost all *Or* transcripts were significantly downregulated (except for *AgOr2*), and additional experiments suggest that it is AgOR2 protein rather than mRNA that mediates this downregulation. Our study shows that ORNs of *Anopheles* mosquitoes (in contrast to *Drosophila*) employ a currently unexplored regulatory mechanism of OR expression, which may be adaptable as a vector-control strategy.

**SIGNIFICANCE STATEMENT:** Studies in *Drosophila melanogaster* suggest that insect Olfactory Receptor Neurons (ORNs) do not contain mechanisms by which Odorant Receptors (ORs) regulate OR expression. This has proved useful in studies where ectopic expression of an OR in *Drosophila* ORNs confers responses to the odorants that activate the newly expressed OR. In experiments in *Anopheles gambiae* mosquitoes, we found that ectopic expression of an OR in most *Anopheles* ORNs dampened the activity of the expressing neurons. RNA-seq analyses demonstrated that ectopic OR expression in *Anopheles* ORNs leads to downregulation of endogenous *Or* transcripts. Additional experiments suggest that this downregulation required ectopic expression of a functional OR protein. These findings reveal that *Anopheles* mosquitoes, in contrast to *Drosophila*, contain a feedback mechanism to regulate OR expression. Mosquito ORNs might employ regulatory mechanisms of OR expression previously thought to occur only in non-insect olfactory systems.

---

Malaria is a major cause of human mortality worldwide (1), and it is a global health imperative to prevent the spread of this disease. Malaria is caused by *Plasmodium* parasites transmitted by the bite of infected *Anopheles* mosquitoes. To date, antimalarial drugs have been the mainstay of control against malaria, and over the past 15 years, these treatments – along with the distribution of insecticide-treated bed nets – have contributed to an overall reduction of disease transmission. However, the eventual eradication of malaria likely rests on a multidisciplinary approach that integrates our knowledge of both host and vector biology (2). For example, impairing the ability of the insect vector to bite a human host may further reduce incidences of infection. As such, disrupting the behaviors that bring mosquitoes to humans could dramatically reduce the prevalence of malaria (3).

Female *Anopheles* mosquitoes locate and approach humans (‘host-seek’) based on specific cues, such as human-derived odors and exhaled CO_2_, moisture and heat emissions, and body shape. The primary way that mosquitoes host-seek is through their sense of smell (olfaction). Mosquitoes have evolved a complex repertoire of chemoreceptors that respond to chemical stimuli such as Ionotropic Receptors (IRs), Gustatory Receptors (GRs), and Odorant Receptors (ORs). Of these, the ORs play a substantial role in mediating how a mosquito responds to human odor (4). ORs form heterotetramer complexes with an obligate co-receptor known as ORCO (5). OR-ORCO complexes are expressed on Olfactory Receptor Neurons (ORNs) of the mosquito’s sensory appendages: the antennae, maxillary palps, and labella (6). ORCO-positive ORNs are housed in ‘sensilla’ or sensory hairs of these appendages. Each sensillum typically contains 1-4 ORNs that each express a unique OR, of which there are 75 in the *Anopheles gambiae* genome (7). These ORNs are classified and named by the OR gene they express, and each ORCO-positive ORN class targets a specific brain region of the mosquito Antennal Lobe (AL) known as a glomerulus (8, 9). The decision to approach a human is a direct result of activated ORCO-positive ORNs targeting a *specific* combination of glomeruli (10) (**Fig 1a,b**).

**Figure 1.**
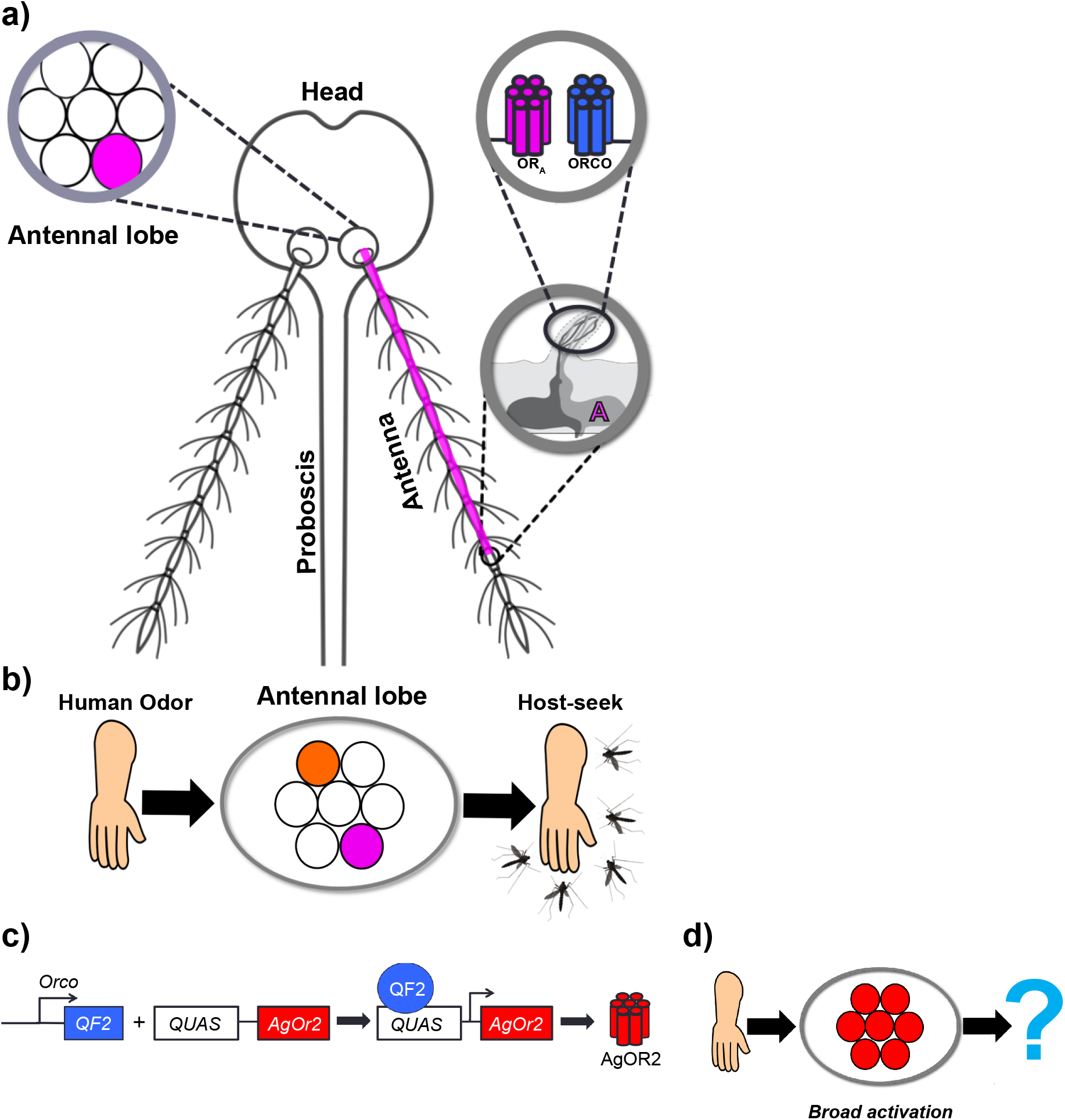
Strategy to manipulate the olfactory system of *Anopheles* mosquitoes. a) Anatomy of odorant receptor guided olfaction. Mosquitoes smell odors in the environment using three olfactory organs: the proboscis, the maxillary palps (not shown), and the antennae. A single antenna is made up of 13 segments called ‘flagellomeres.’ Each flagellomere is covered with sensory hairs called ‘sensilla,’ a single one of which houses up to 4 olfactory neurons. An olfactory neuron expresses 1 of 3 chemosensory gene families: the *ionotropic receptors*, the *gustatory receptors*, or the *odorant receptors*. The *odorant receptor* gene family plays an important role in host-seeking behavior. There are 75 different ORs in the *Anopheles gambiae* genome, each of which is sensitive to specific odors in the environment. Only 1 OR (OR_A_) is expressed *per* olfactory receptor neuron (ORN, labeled ‘A’). At the dendrites of ORN A, OR_A_ couples with the obligate co-receptor, ORCO. When odor binds to OR_A_-ORCO complexes, ORN A becomes active and sends its excitatory signal down its axon, targeting a discrete brain region of the mosquito antennae lobe (AL) called a ‘glomerulus’ (shown as a pink circle). **b) A human-specific odor code in the mosquito brain**. Human odors bind to specific ORs, activating the ORNs on which they are expressed. Activated ORNs target discrete glomeruli (pink, orange) in the AL to guide host-seeking. **c) Olfactogenetics strategy**. Using the Q-system, the *Orco-QF2* transgene (6) was combined with the effector construct, *QUAS-AgOr2*. The combination of these transgenes causes AgOR2 to be expressed in *Orco*-positive ORNs. **d) Test of the olfactogenetics strategy**. AgOR2 responds to major components of human odor such as indole and benzaldehyde. We hypothesized that *Orco>AgOr2* mosquitoes would experience broad activation of the majority of the AL in the presence a human, thereby dysregulating the human-specific odor code in the mosquito brain (**Fig. 1b**). *Orco>AgOr2* mosquitoes would then be evaluated on whether they showed reduced host-seeking behavior.

Disrupting this specificity by activating all ORCO-positive ORNs in the presence of human odor has been hypothesized to prevent host-seeking (11, 12). Studies in *Drosophila*, an insect model of olfaction, show that binary systems can be used to express non-native ORs in ORCO-positive ORNs to confer ectopic neuronal responses without interfering with native olfactory physiology. Therefore, we examined whether this strategy could be used to disrupt host-seeking in *Anopheles gambiae* mosquitoes. To test this, we expressed the *Anopheles gambiae odorant receptor 2 (AgOr2)* in all *Orco*-positive ORNs. AgOR2 is highly attuned to major components of human odor such as benzaldehyde and indole (13), and in the presence of human odor, all *Orco*-positive neurons expressing AgOR2 should become active. Surprisingly, when we evaluated the olfactory physiology of these experimental mosquitoes, we found that they exhibited reduced responses not only to AgOR2’s cognate ligands (benzaldehyde and indole) but to odors in general. To investigate the molecular basis of this phenotype we looked for signatures of disfunction at the level of the transcriptome. Using RNA-seq to compare transcript levels from wild-type antennae to those ectopically expressing *AgOr2*, we discovered that *odorant receptor* isoforms were significantly downregulated in the experimental line while the remaining transcripts were largely unchanged. Additional experiments revealed that it is AgOR2 protein rather than *AgOr2 mRNA* that reduces native *odorant receptors* levels. Overall, our study suggests the existence of a feedback mechanism of *odorant receptor* regulation whereby an OR protein downregulates the transcripts of alternative *Or* genes.

## RESULTS

### Ectopically Expressing *AgOr2* in *Orco*-positive Neurons Impairs Olfactory Physiology

To activate *Orco*-positive ORNs in the presence of human odor, we used ‘olfactogenetics,’ a technique whereby a specific volatile odorant is used to activate a defined set of OR-expressing neurons (14). To accomplish this, an *Or* with known response properties is ectopically expressed in ORNs of interest through the use of a binary expression system, such as the Q-system (15) or the *UAS*-Gal4 system. Thus, in the presence of odors in which the introduced OR normally responds, olfactory receptor neurons with ectopic expression become active.

In 2016, the Q-system was introduced into *Anopheles*, making it possible to adapt olfactogenetics in mosquitoes (6). As a binary expression system, the Q-system works by directing the expression of a specific gene into a specific cell population. This particular system relies on two elements: QF2 and *QUAS*. The QF2 transcription factor is expressed under the control of a cell-type specific enhancer/promoter and binds to its upstream activating sequence, *QUAS*. Once bound by QF2, *QUAS* initiates transcription of its effector gene. To ectopically activate *Orco*-positive ORNs in the presence of human odor, we combined a mosquito line containing *Orco-QF2* (6), which contains a fusion between the presumptive enhancer and promoter regions of the gene *Orco* and the transcription factor QF2, with an effector line containing a *QUAS* transgene upstream of the *Anopheles gambiae odorant receptor 2, AgOr2 (QUAS-AgOr2)*. Thus, experimental animals exhibit ectopic expression of *AgOr2* in all *Orco*-positive ORNs (**Fig. 1c**).

AgOR2 is highly attuned to major components of human odor such as benzaldehyde and indole (16) and it was expected that *Orco*-positive ORNs of *Orco>AgOr2* mosquitoes would become active in the presence of these cognate ligands, essentially activating the majority of the olfactory system during host-seeking (**Fig. 1d**). To test the functional activity of olfactory receptor neurons ectopically expressing *AgOr2* in mosquitoes, we imaged the calcium response of *Orco>AgOr2* antennal segments in a *QUAS-GCaMP6f* background (*Orco>AgOr2,GCaMP6f*). Surprisingly, *Orco>AgOr2,GCaMP6f* antennae showed a dampened response not only to various concentrations of benzaldehyde and indole, but to odors in general (**Fig. 2a**). For example, octenol potentially activates 31 different *Anopheles gambiae* ORs (16), but the olfactory receptor neurons of the experimental mosquitoes did not show a response to this odor. In *Drosophila*, a single ORN class can drive behavior at spike rates as low as 10-20Hz (10). When we stimulated *Orco>AgOr2,GCaMP6f* with 5 additional odors known to activate 14-16 different ORN classes at rates higher than 50 spikes/sec (16), the olfactory receptor neurons still did not exhibit odor-induced responses (**Fig. S1**).

**Figure 2.**
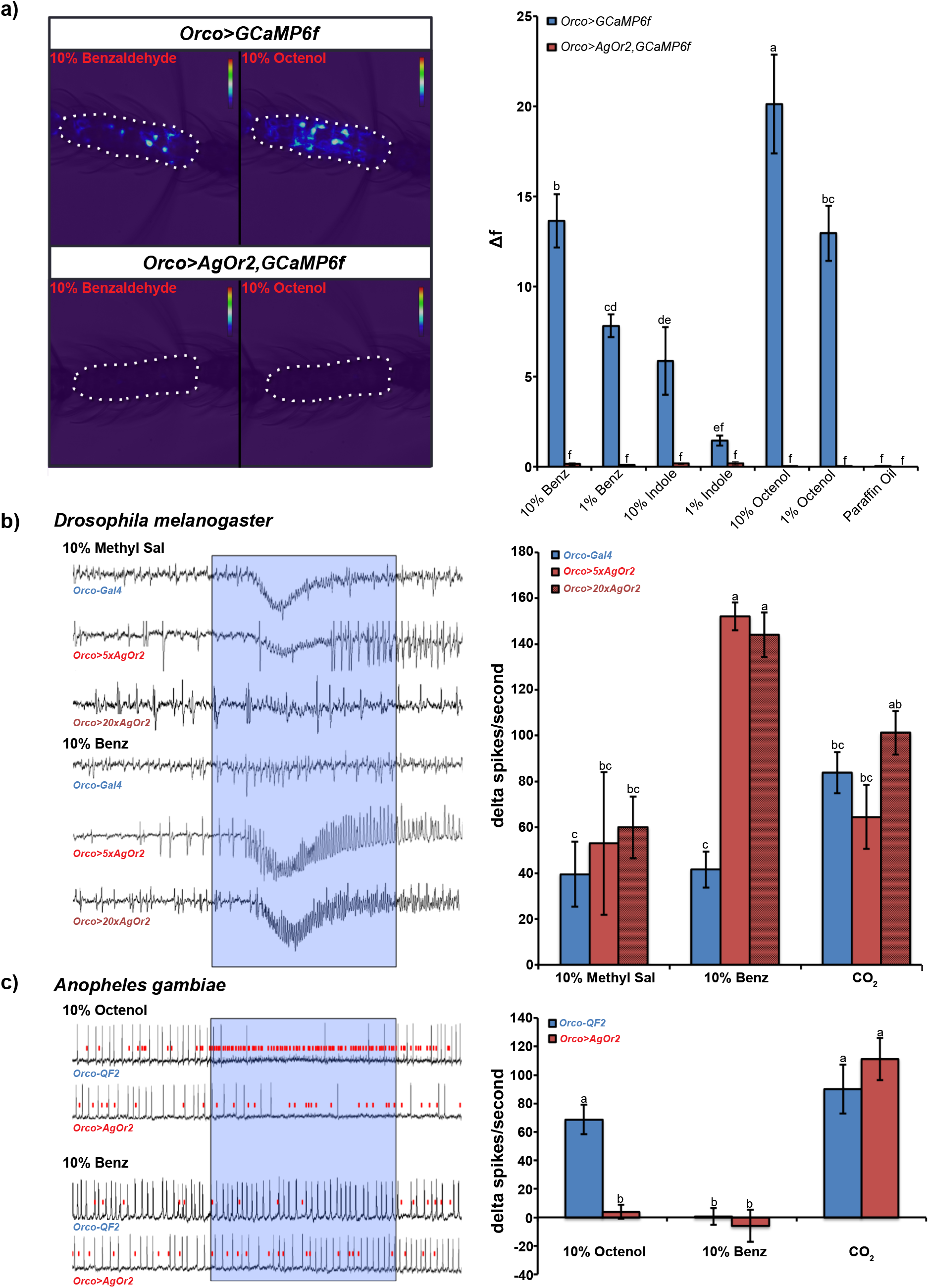
Olfactogenetics impairs *Anopheles* but not *Drosophila Orco*-positive ORNs. **a) *Orco>AgOr2,GCamp6f* mosquitoes show impaired olfactory responses to human odorants**. The activity of olfactory receptor neurons in antennal segment 11 (outlined with a white dotted line) was detected by calcium imaging of *Orco*-positive neurons expressing *GCaMP6f* (51). Relative to controls (*Orco>GCaMP6f*), mosquito antennal segments with ectopic expression of *AgOr2* and *GCaMP6f (Orco>AgOr2,GCaMP6f*) show impaired responses to benzaldehyde (10% and 1%) and indole (10%), the cognate ligands of AgOR2. They also show dampened responses to octenol (10% and 1%). A two-way repeated measures ANOVA was conducted to test the effect of odor and genotype on calcium responses. We found a significant effect at the *p*<0.0001 level. Groups with different letter values (a-f) are statistically different as determined by the Tukey post hoc HSD test. Each sample included in the analysis was taken from a different female mosquito. n_*Orco>GCaMP6f*_ = 9, n_*Orco>AgOr2,GCaMP6f*_ = 8. **b) Ectopic expression of *AgOr2* in *Drosophila* ab1 sensilla does not impair ORN physiology**. The cognate ligand of OR10a-expressing neurons is methyl salicylate (Methyl Sal). Driving *AgOr2* into this neuronal group using the *5xUAS* or *20xUAS* effector lines does not affect OR10a’s response to Methyl Sal. *Orco*-positive ORNs of *Orco>5xAgOr2* and *Orco>20xAgOr2* animals show an ectopic response to benzaldehyde (Benz). The presence of the *Orco*-negative CO_2_ neuron was used to verify that recordings were taken from the ab1 sensillum. The activity of the CO_2_ neuron (trace not shown) is not affected by the experimental manipulation. Odor or CO_2_ stimulus was delivered in the timeframe denoted by the blue translucent box. A two-way repeated measures ANOVA was used to determine significance of genotype and odor on delta spikes/second at the *p<0.0001* level. Groups with different letter values (a-c) are statistically different as determined by the Tukey post hoc HSD test. 2-3 Females *per* genotype were analyzed. The number of sensilla evaluated for each group: n_*Orco-GAL4*_ = 9; n_*Orco>5xAgOr2*_ = 4; n_*Orco>20xAgOr2*_ = 5. **c) Olfactogenetics impairs *Orco*-positive ORN physiology in *Anopheles***. Single sensillum recordings from the *Anopheles* maxillary palp capitate peg sensilla. The cognate ligand of OR8-expressing neurons is octenol. Driving *AgOr2* into this neuron group interferes with OR8’s response to octenol (smallest spiking neurons, red lines indicate OR8 activity). The ORNs ectopically expressing *AgOr2* do not respond to benzaldehyde (Benz). The presence of the *Orco*-negative CO_2_ neuron was used to verify that recordings were taken from a cp sensillum. The activity of the CO_2_ neuron (trace not shown) was not affected by the experimental manipulation. Odor or CO_2_ was delivered in the timeframe denoted by the blue translucent box. A two-way repeated measures ANOVA was used to determine that there was a significant effect at the *p*<0.005 level of odor and genotype on delta spikes/second. Groups with different letter values (a-b) are statistically different as determined by the Tukey post doc HSD test. 2-3 Females *per* genotype were analyzed. The number of sensilla evaluated for each group: n*_Orco-QF2_* = 6; n*_Orco>AgOr2_* = 5. Error bars represent the standard error (SEM).

One possibility for the observed olfactory defects (**Fig. 2a, Fig. S1**) is that the *QUAS-AgOr2* transgene inserted into an endogenous olfactory gene and disrupted olfactory functions. As determined by splinkerette mapping (17), the insertion site of the *QUAS-AgOr2* line examined in **Fig. 2a and Fig. S1** (line 1) is located in an intergenic region on chromosome 3R between the genes ACOM029303 and ACOM029196. While it is unlikely that this particular insertion site would disrupt the function of an olfactory gene necessary for ORN physiology, we extended our studies to analyze the physiology of two additional *QUAS-AgOr2* lines inserted into different regions of the genome. We established line *QUAS-AgOR2#2*, which maps to an intergenic region between ACOM036217 and ACOM036230, and line *QUAS-AgOR2#*3, which could not be mapped by splinkerette PCR. When driven into *Orco*-positive cells, all three lines show similar defects in olfactory physiology when compared to wild-type (**Fig. S2**). Furthermore, since these lines were tested as heterozygotes, any recessive mutation in an olfactory gene caused by the *QUAS-AgOr2* insertion should be compensated by the wild-type allele. Overall, these data show that the dominant negative olfactory phenotype (**Fig. 2a, Fig. S1, Fig. S2**) is a consequence of ectopic *AgOr2* expression rather than the genomic insertion site of the *QUAS-AgOr2* element.

### Olfactogenetics Impairs *Anopheles* but not *Drosophila Orco*-positive ORNs

Ectopically expressing an *Or* in *Drosophila* ORNs causes the expressing neuron to activate in the presence of the introduced OR’s odor ligand (14, 16, 18–20). One possibility as to why *Orco>AgOr2* cells in the mosquito did not respond to benzaldehyde or indole was because the *AgOr2* sequence used to create the transgenic *QUAS-AgOr2* line was perhaps acting in a dominant-negative manner to disrupt ORCO/OR_X_ ion channels. To test this, we ectopically expressed *AgOr2* in *Drosophila Orco*-positive neurons using the GAL4-UAS system and measured the response rate of neurons housed in the ab1 sensilla using Single Sensillum Recordings (SSR). Ab1 contains 4 olfactory neurons: 3 of which are *Orco*-positive and express *Or10a, Or42b*, and *Or92a*, and one of which is *Orco*-negative and expresses the gustatory receptor *Gr21*, which responds to CO_2_. To determine whether ectopic expression of *AgOr2* impairs native OR responses to their cognate ligands, responses of the OR10a-expressing neuron to methyl salicylate were measured. We found no difference in how control (*Orco-GAL4*) and experimental (*Orco>5xAgOr2*) sensilla responded to methyl salicylate, indicating that *AgOr2* expression does not interfere with the olfactory physiology of the neuron in which it is expressed. Furthermore, native responses of OR10a were not affected when an even higher dosage of *AgOr2* was driven into the neuron (*Orco>20xAgOr2*) (**Fig. 2b**). When we puffed benzaldehyde over the experimental preparation, *Orco*-positive cells in the *Drosophila* ab1 sensilla ectopically respond. Interestingly, sensilla that have higher levels of ectopic *AgOr2* can still maintain ectopic responses without compromising neuron function. When we used the stronger effector line (*20XUAS*) to ectopically express *AgOr2* in *Drosophila Orco*-positive neurons, the olfactogenetics approach continued to work and olfactory physiology was not impaired.

The *Anopheles* capitate peg (cp) sensillum is similar to the ab1 sensillum of *Drosophila* as it contains *Orco*-positive ORNs, which express *Or8* and *Or28*, and one *Orco*-independent olfactory neuron that responds to CO_2_. When we ectopically expressed *AgOr2* in *Orco*-positive ORNs in the mosquito and recorded from cp sensilla, we found that the OR8-expressing neuron’s response to octenol, its cognate ligand, was eliminated (**Fig. 2c**). In addition, neither OR8 nor OR28-expressing neurons ectopically respond to benzaldehyde. Similar to *Drosophila*, the genetic manipulation does not affect the physiology of the *Orco*-negative CO_2_-responsive neuron (**Fig. 2c**). These data indicate that olfactogenetics affects the physiology of *Drosophila* and *Anopheles Orco*-positive ORNs differently: in flies, ectopically expressing *Or*s does not affect endogenous neuronal function, whereas in *Anopheles* mosquitoes, ectopic expression of an *Or* disrupts the function of the olfactory receptor neurons.

### Ectopic AgOR2 protein eliminates olfactory responses in *Orco*-positive cells

It is possible that driving an *Or* into an *Anopheles* ORN is cytotoxic, especially if the neuron experienced continual stimulation from the environment. To assess whether this was the case, we examined the anatomy of the AL and ORN projections in *Orco>AgOr2* mosquitoes, to check for intact neuronal processes. As shown previously (6), ablating *Orco*-positive cells would cause their ORN projections to be eliminated in the AL. Using immunochemistry to visualize ORN terminals in the AL, both the ORN projections and the AL in general remained intact, indicating that the lack of ORN responses to odor was not due to the death and elimination of neurons ectopically expressing *AgOr2* (**Fig. S3**). Next we tested if driving any generic transmembrane protein into *Anopheles* ORNs also hindered olfactory physiology. To evaluate this, we used the *Orco-QF2* driver to ectopically express the transmembrane protein *mCD8::GFP* (6) into all *Orco*-positive neurons. There were no differences in the odor-induced responses between control and *mCD8::GFP-positive* ORNs (**Fig. S4**), suggesting that ectopic expression of another transmembrane protein did not silence ORN activities.

Ectopic *AgOr2* expression may inhibit olfactory responses either at the level of the *AgOr2* RNA or at the level of AgOR2 protein. To distinguish between these two possibilities, we created a transgenic mosquito line containing a mutated version of *AgOr2 (mutAgOr2)* that contained a mutation in the start codon of *AgOr2* such that *QUAS-mutAgOr2* produced mRNA that cannot be translated when combined with *Orco-QF2*. We also induced a frameshift mutation at a second in-frame ATG site in *mutAgOr2* to eliminate the possibility of having an alternative open reading frame used during translation. We crossed *QUAS-mutAgOr2* with *Orco-QF2,QUAS-GCaMP6f* and found that the calcium responses were not compromised (**Fig. 3**); only when the functional version of the protein was expressed (*QUAS-AgOr2*) was the odor response impaired (**Fig. 2a,c, Fig. S1, Fig. S2**). Taken together, these data indicate that AgOR2 protein itself, and not mRNA, was largely responsible for the olfactory defect of *Orco*-positive cells.

**Figure 3.**
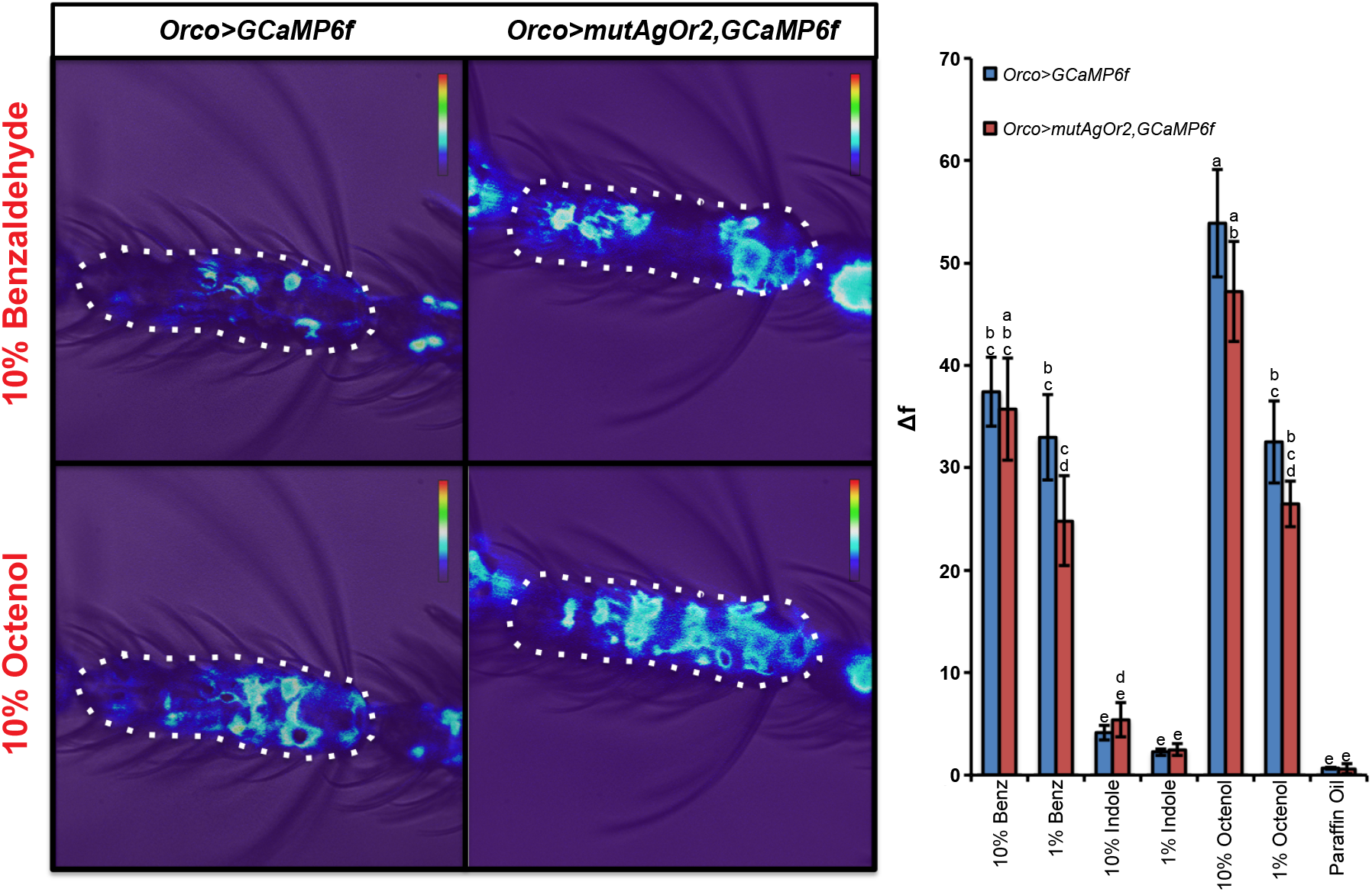
AgOR2 protein is required for the dominant negative olfactory phenotype caused by *Orco>AgOr2* expression. The *mutAgOR2* transgene contains an introduced point mutation in the start codon of *AgOr2* and a frameshift mutation at a second in-frame ATG site of the gene. Odor-evoked responses (Δf) were calculated from the 11^th^ segment of the mosquito antennae (outlined by dotted lines). Representative control and experimental calcium imaging responses to 10% benzaldehyde and 10% octenol are shown. Antennae from *Orco>mutAgOr2,GCaMP6f* mosquitoes show no difference in responses to odors from control (*Orco>GCaMP6f*). A two-way repeated measures ANOVA was used to determine that there was a significant effect at the *p*<0.005 level of odor and genotype on calcium responses. Groups with different letter values (a-d) are statistically different as determined by the Tukey post hoc HSD test. Error bars represent the standard error (SEM). Each sample included in the analysis was taken from a different female mosquito. n*_Orco>GCaMP6f_* = 11, n_*Orco>mutAgOr2,GCaMP6f*_ = 5.

### Ectopic AgOR2 Reduces the Transcripts of Native *Ors*

How might AgOR2 protein impair olfactory responses? We hypothesized this may occur as a result of 1) regulatory mechanisms affecting *Or* transcription, stability, and/or degradation rates or 2) regulatory mechanisms and/or defects in OR protein function. These transcriptional or post-translational mechanisms may act together, or independently. To distinguish between these two possibilities, we performed isoform-level RNA-seq on antennae isolated from control and *Orco>AgOr2* samples. Of the ~13,000 transcript isoforms detected, only 83 were differentially expressed in the *Orco>AgOr2* mosquito antennae. Interestingly, half of these differentially expressed isoforms (41/83) were *Ors*, which were all downregulated in *Orco>AgOr2* antennae, except for *AgOr2*, which was highly upregulated (**Fig. 4a**). As shown by the *Or* isoform comparison heatmaps in **Figure 4b**, control *Or* levels are higher than those in *Orco>AgOr2*, with the exception of *AgOr2*, which is upregulated.

**Figure 4.**
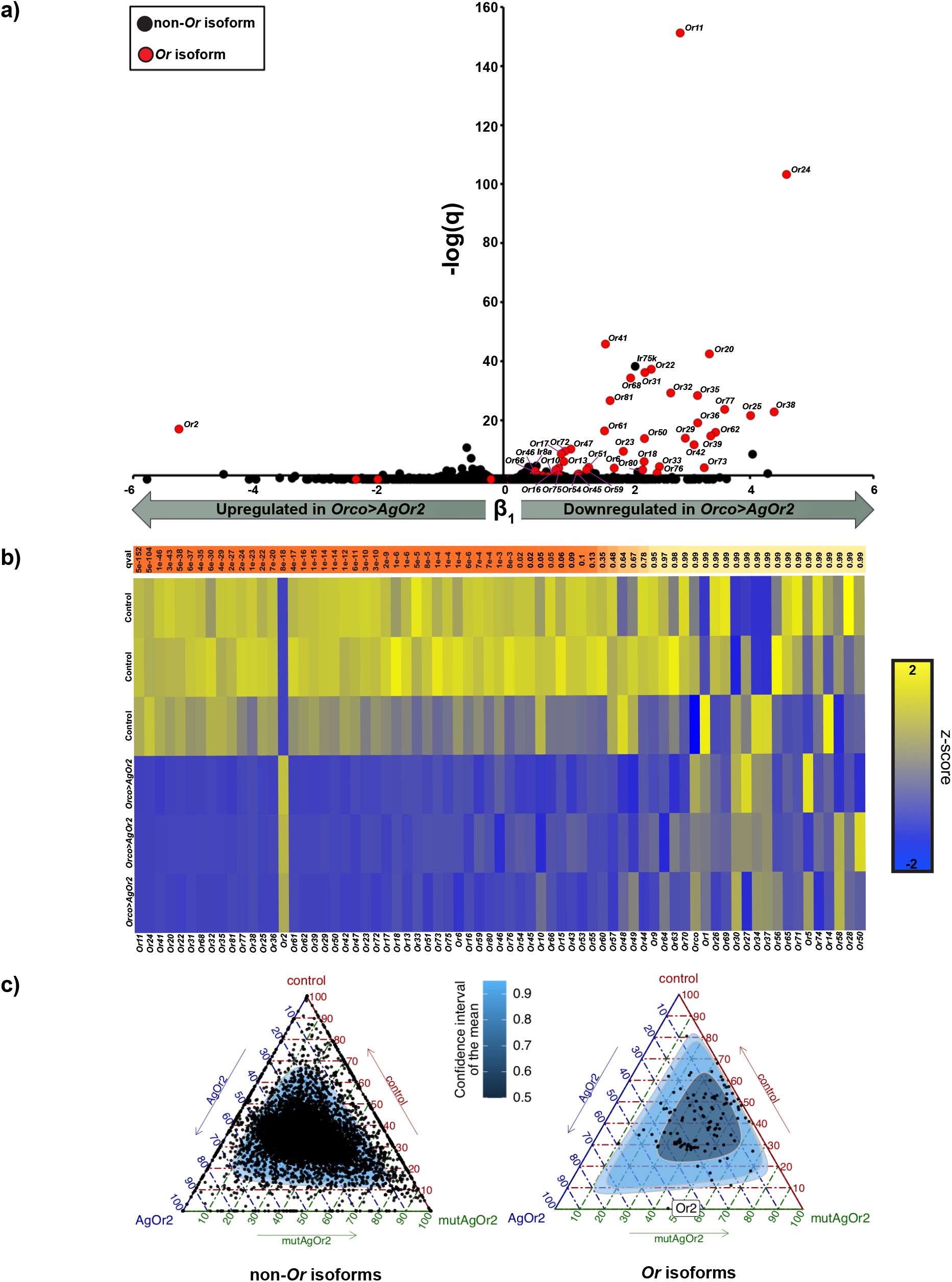
Ectopic AgOR2 protein reduces the transcript levels of *Or* isoforms. a) Volcano plot of differentially expressed isoforms. Using Wald tests, we evaluated whether the 13224 isoforms present in 3 triplicates of control and 3 triplicates of *Orco>AgOr2* antennae (~200 antennae *per* sample) were differentially expressed. *Non-odorant receptor* and *odorant receptor* isoforms are shown as black or red dots, respectively. Only 0.63% of the transcriptome is differentially regulated in *Orco>AgOr2*, where 49% of those transcripts are *Ors*. −log(q) is the level of significance of β_1_, which, for each isoform, is defined as TPM_control_ - TPM_experimental_. The 41 *Ors* found significant from the Wald tests are labeled according to their gene annotation in the volcano plot. All *Ors* (with the exception of *AgOr2*) that are differentially expressed are downregulated in the experimental condition. Interestingly, *Ir8a* and *Ir75k* (indicated on the volcano plot) are downregulated in *Orco>AgOr2*. The remaining *Irs* and *Grs* are unaffected. **b) Heatmap of the *Or* gene family**. A z-score was computed for each cell in the heatmap by subtracting the mean isoform TPM from the cell’s TPM divided by the standard deviation of the isoform TPM. *Ors* are sorted along the X-axis according to their significance level (q value) from the Wald tests in **a**. Darker orange is most significant, yellow is not significant. *Ors* are downregulated in *Orco>AgOr2* samples, with the exception of *AgOr2*, which is upregulated. **c) Ectopic AgOR2 protein is required for the observed downregulation of native *Or* transcripts**. Ternary plots were used to visualize the relative ratio of a genotype to an isoform’s relative abundance level using the formula: (TPM_genotypeX_)/(mean TPM_control_ +mean TPM_Orco>AgOr2_+mean TPM_Orco>mutAgOr2_)*100. (left) There are equal ratios of transcript abundance levels for non-*Or* genes among the three genotypes (right). However, the relative contribution to transcript abundance levels of control, *Orco>AgOr2*, and *Orco>mutAgOr2* are skewed in the *Or* gene family such that *Or* gene levels in *Orco>mutAgOr2* and control are relatively similar and higher than *Orco>AgOr2* levels, with the exception of *AgOr2* itself (whose abundance is similar between *Orco>AgOr2* and *Orco>mutAgOr2*).

If AgOR2 ectopic protein was responsible for modulating the steady state abundance of native *Or* transcripts, then the relative abundance of the *odorant receptor* gene family isoforms in *Orco>mutAgOr2* antennae should not be affected. To test this, we extracted biological triplicates of mosquito antennae from female mosquitoes of the *Orco>mutAgOr2* line and compared the isoform abundance levels of this group to the original RNA-seq dataset. Ternary plots depicting the relative abundance of isoform expression levels in control, *Orco>mutAgOr2*, and *Orco>AgOr2* conditions as a position on an equilateral triangle were used to explore how *Or* and other gene sets were differentially expressed. To depict how control, *Orco>mutAgOr2*, and *Orco>AgOr2* contribute to relative isoform abundances of non-*Or* genes and *Or* genes, we created 2 discrete ternary plots. As expected, the relative contribution of control, *Orco>mutAgOr2*, and *Orco>AgOr2* for all non-*odorant receptor* isoform in the transcriptome was roughly equal (34.3%:33%:32.6%). However, when removing the contribution of the ectopically induced *AgOr2* from the *Or* isoform pool, the ratio for the *odorant receptor* family was skewed to 45.8%:37.8%:16.4% (control:*Orco>mutAgOr2:Orco>AgOr2*) (**Fig. 4c**), indicating that the relative contribution of an *Or’s* abundance level was roughly equal among control and *Orco>mutAgOr2* groups, but reduced in *Orco>AgOr2*. Results of the gene set enrichment test and pairwise comparisons of *Or* isoforms can be viewed in **Figure S5**. The majority of native *Ors* are downregulated in *Orco>AgOr2* but not in *Orco>mutAgOr2*, with four exceptions: *Or2, Or16, Or17*, and *Or33*.

### As visualized by the onset of ORCO expression, ectopic AgOR2 is driven immediately after larval-pupal ecdysis

We next examined when, during the life cycle of the mosquito, the downregulation of native *Or* genes via ectopic AgOR2 expression might occur. Since ectopic AgOR2 expression is dictated by the *Orco* enhancer/promoter region (**Fig. 1c**) (6), AgOR2-induced olfactory silencing of native *Or* genes should coincide with the ORCO expression pattern. Mosquitoes experience four different developmental stages during their lifespan: egg, larvae, pupae, and adult stages. During the pupal stage, adult features of the mosquito olfactory system take shape (21, 22), so we examined this stage of development for the onset of ORCO expression. To document this, we extracted and stained pupal antennae from *Orco>mCD8::GFP* mosquitoes every 4hrs starting at the onset of pupal development (After Puparium Formation: 0-2hr APF) to just before eclosion (20-22hr APF), which happens at 24hrs APF. ORCO (as reported by mCD8) was expressed immediately after pupal ecdysis, just 0-2hrs APF (**Fig. S6a**). Interestingly, ORCO expression turns on gradually in the developing antennal ORNs, where each flagellomere gains ~15 Orco-positive cells every 4hrs, starting from ~8 cells at 0-2hr APF and ending with ~87 cells per flagellomere at 4hrs before eclosion (**Fig. S6b**). These data suggest that ectopic expression of AgOR2 (which should coincide with ORCO) also occurs throughout pupal development, and might impinge upon developmental mechanisms that regulate *Or* expression. The timepoint at which native *Or* genes begin to express in *Anopheles* is not currently known, but in *Aedes aegypti* several *Ors* (in addition to *Orco*, as detected by *in situs*) are present 3/4ths of the way through the pupal stage (23). This suggests that in natural conditions, negative inhibition of *AgOr* genes by AgOR protein might occur during the pupal stage, when adult features of the olfactory system are being developed.

### Impairing *Orco*-positive neuron responses does not inhibit host-seeking behavior in *Anopheles* mosquitoes

In the absence of an *Orco* mutant, the olfactory phenotype of *Orco>AgOr2* mosquitoes presented the opportunity to test whether *Anopheles* mosquitoes without ORN function still host-seek. DeGennaro et al. 2013 (4) found that a null mutation of the *Orco* gene in *Aedes* mosquitoes did not prevent host-seeking behavior (presumably due to redundant function of IR neurons (24)), but whether the malaria mosquito would behave similarly is unknown. To test whether *Orco>AgOr2* mosquitoes showed reduced attraction towards human hosts, we used the ‘host-proximity assay’ (4), a population assay that measures the proportion of females that come into olfactory (but not physical) contact with a human arm. As found with *Orco* mutant *Aedes* mosquitoes (4), *Anopheles* mosquitoes without functional *Orco*-positive neurons were still attracted to a human host (**Fig. 5a-c)**.

**Figure 5.**
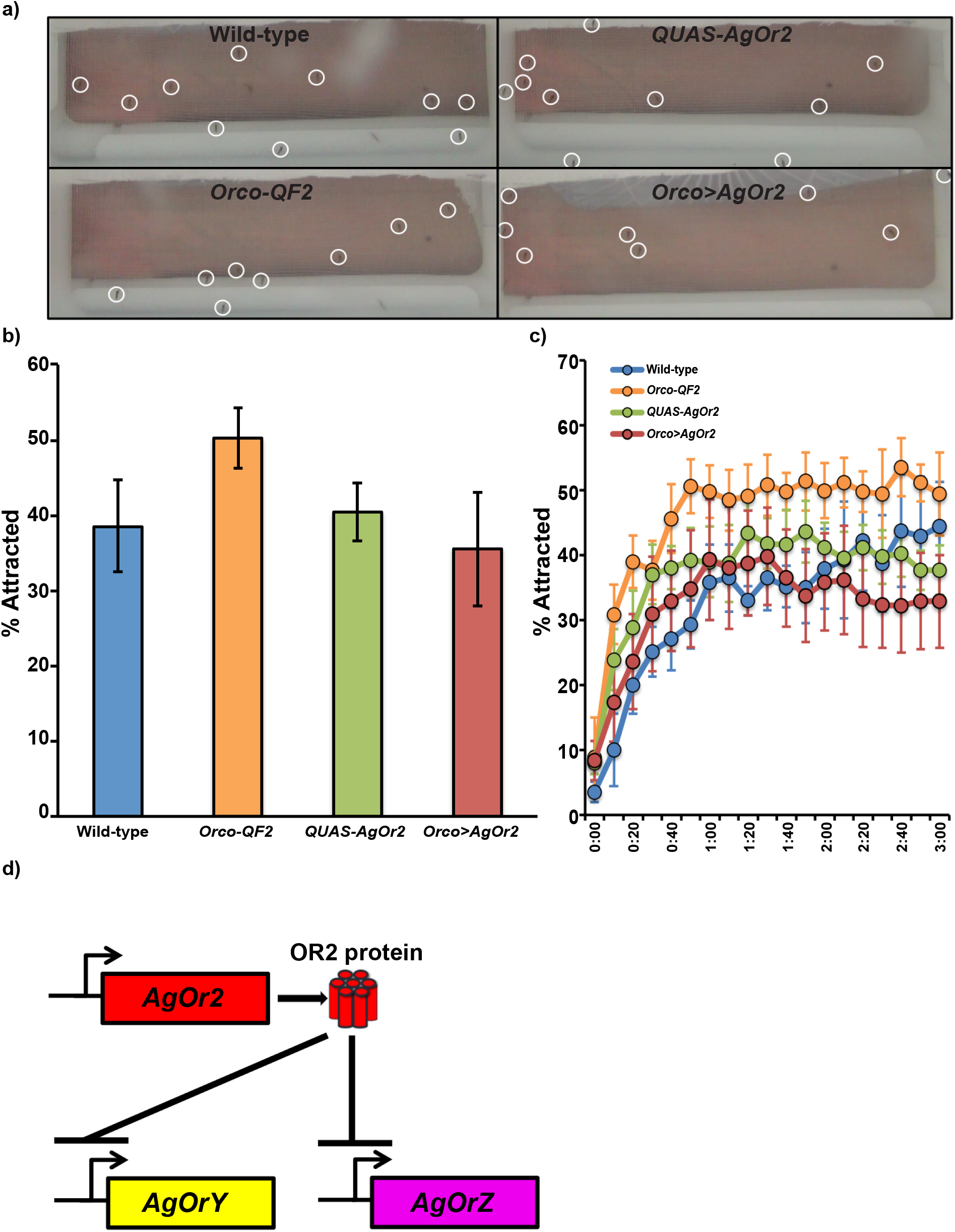
*Orco>AgOr2* mosquitoes remain attracted to a human host. **a)** Representative images of wild-type, *Orco-QF2, QUAS-AgOr2*, and *Orco>AgOr2* mosquitoes (circled) in the host-proximity assay. Mosquitoes attracted to an arm (2.5cm from the cage) that land on the net are counted. **b) Results of the host-proximity assay**. A one-way ANOVA between subjects was conducted to compare the effect of genotype on % attraction. There was no effect of genotype on % attraction at the *p*<0.05 level for the four groups (*F(3)* = 1.08, *p*=.37). Error bars represent the standard error (SEM). 20-30 female mosquitoes were tested *per* trial. The number of trials *per* genotype: n_Wild-type_ = 5; n_*Orco-QF2*_ = 5; n_*QUAS-AgOr2*_ = 7; n_*Orco>AgOr2*_ = 7 **c) Time course of mosquito attraction towards a human host by genotype**. Over the course of 3 minutes, there was no difference in the % of mosquitoes attracted to a human host. **d) Summary Model: ectopic AgOR2 negatively regulates the expression of *Or* transcripts**. Our data implicates a mechanism of negative regulation of most *Or* transcripts (for example, *OrY* and *OrZ*) by ectopic AgOR2 protein.

## DISCUSSION

We report for the first time, to our knowledge, an unexplored mechanism of *Or* regulation in *Anopheles* mosquitoes that diverges from the dogma established in *Drosophila*. A main finding of our paper – that <u>OR protein</u> downregulates the expression of native *Ors* (**Fig. 5d**) – is a pattern more closely resembling OR-regulatory processes in mice than those of *Drosophila*. As this (**Fig. 2b**) and previous studies have demonstrated (14, 18, 20) driving ectopic expression of an *Or* in non-native *Orco*-positive ORNs does not disrupt *Drosophila* neuron function whereas in *Anopheles* it does (**Fig. 2a,c, Fig. S1, Fig. S2**). Furthermore, ectopically expressing an *Or* gene in *Drosophila* ORNs does not affect gene expression of native *Ors* (20), whereas a similar manipulation in *Anopheles* leads to robust changes in gene expression (**Fig. 4**).

From these experiments, we hypothesize unexpected similarities of *Or* gene regulation between *Anopheles* and mice. For example, both *Anopheles* and mice contain a negative feedback loop by which OR protein inhibits the expression of alternative *Ors*. In *Anopheles* mosquitoes (**Fig. 3, Fig. 4**) as was the case in mice, this feedback pathway requires intact OR protein (with 3 exceptions: see **Fig. S5**) since expressing mutant *Or* genes lacking either the entire coding sequence or the start codon permits a second *Or* gene to be expressed (25–28). Frameshift mutations also allow for the co-expression of functional *Or* genes (25). It remains to be determined if this repression in *Anopheles* olfactory receptor neurons utilizes similar cellular machinery as in mice (29).

What might be the selective advantage of an *Anopheles* mosquito *Or* feedback mechanism? One possibility is that it increases the likelihood that a single olfactory receptor neuron expresses only 1 OR. This is an important developmental mechanism in mice, where ORs play an instructive role in guiding olfactory receptor neuron axonal projections to the olfactory region of the brain (30–32). In analogous experiments to those presented here, when multiple ORs are genetically engineered to co-express in a single olfactory receptor neuron in mice, the topographic map of projections to the olfactory center in the brain is perturbed (33). While we do not know when native *Or* genes turn on or when ORNs target the AL in *Anopheles*, our methods expressed *AgOr2* in *Orco* neurons as early as 0-2hr APF (**Fig. S6**). If ORNs have not yet targeted the AL at this early stage, our data suggests that *Ors* in mosquitoes are likely not involved in axon guidance, as we did not detect major deformations of ORN targeting or AL structure in the adult brain (**Fig. S3**). What might be another biological process influenced by OR gene regulation? If this mechanism functions during adulthood, it could be important for synchronizing a mosquito’s dynamic biological needs in the environment with the physiology of olfactory receptor neurons. Mosquitoes rely heavily on their sense of olfaction to integrate ecologically relevant stimuli that change over the course of their adult lifespan. When females first eclose, they are uninterested in host odor (34) and instead actively search for sugar-rich resources from plants to supplement their nutrient reserves. After a period of ~4 days post eclosion, they develop an attraction to host odors (34, 35) and following a bloodmeal will experience a refractory period to host odor until after oviposition. These changes in behavior have been correlated with changes in chemosensory gene transcript abundance (34, 36). It would be interesting to determine if expression of alternative *Ors* in ORNs only during adulthood – when endogenous ORs are already chosen – can alter *Or* gene expression, or if precocious expression of *Ors* takes on an ‘early-bird-gets-the-worm’ paradigm (37).

*AgOrs* can be co-expressed within the same ORN when transcribed as polycistronic mRNA (38). Polycistronic *Or* mRNA is observed in cases when *Or* genes are clustered tightly together within the genome. Such clustering of *Or* genes is commonplace in mosquito species such as *Anopheles gambiae* (39) and *Aedes aegypti* (40), as well as in mice (41, 42); however, to our knowledge, polycistronic *Or* mRNA has not been observed in rodent olfactory systems. It is possible that polycistronic *Or* expression avoids the negative feedback mechanism of *Or* regulation, enabling the neuron to co-express multiple *Ors*.

While ectopically expressing *AgOr2* downregulates native *Or* genes, it was surprising that *Orco>AgOr2* neurons did not show responses to the cognate ligands of AgOR2 (benzaldehyde and indole). Our RNA-seq data indicates that while native *Or* transcripts are reduced, *AgOr2* transcripts are 3x higher than in wild-type conditions: if *AgOr2* transcripts are being translated, this could lead to 3x the amount of AgOR2 protein. We hypothesize that elevated AgOR2 protein levels may be disrupting the stoichiometry of the AgOR2-ORCO complexes, rendering them non-functional. While research has shown that ORs form stable heteromeric complexes with ORCO (43–46), the stoichiometry underlying OR-ORCO channels is unknown. RNA-seq data from this study and from two independent studies show that in wild-type conditions, there is a conserved ~1:1 relationship between *Orco* and total *Or* transcripts (47, 48). Interestingly, we see that ectopic *AgOr2* expression does not change *Orco* expression (**Fig. 4b**), skewing the ratio in experimental conditions to 1:3. We predict this skewed OR expression in a neuron causes major disruptions at the level of the AgOR2:ORCO protein complex. This may result in a trafficking defect in which AgOR2:ORCO complexes do not migrate to the cell surface. Since Orco is required to traffic OR:ORCO complexes to the dendritic surfaces (43), increasing AgOR2 expression might interfere with this process. Alternatively, changes to the OR:ORCO stoichiometry might result in a functional defect in the channel itself, whereby the malformed complexes cannot respond to odors even if they have been successfully trafficked to the membrane.

This study adds to the mounting evidence that the disruption of a single sensory modality is insufficient to completely eliminate host-seeking behavior in mosquitoes. As first demonstrated in *Aedes aegypti*, mosquitoes with a mutation in the *Orco* gene remain attracted to humans (4); similarly, we find that *Anopheles* mosquitoes with impaired ORCO neuronal function continue to host-seek (**Fig. 5a-c**). In addition to odors, mosquitos are attracted to a wide variety of human-derived cues, including heat, CO_2_, visual stimuli, and moisture; and so one sensory modality is likely able to compensate for the loss of another (4, 24, 49, 50).

Our study uncovers the existence of a mechanism of *Or* regulation in insects whereby expression of an OR protein results in the downregulation of other native *Or* gene isoforms (**Fig. 5d**). This work lays the foundation to explore specific cellular mechanisms utilized by mosquito olfactory receptor neurons to regulate *Or* expression. A mechanism of OR regulation in mosquitoes may also be a target for vector-control strategies to alleviate the spread of vector-borne diseases.

## METHODS

### Insect stocks

#### Mosquitoes

*Orco-QF2* and *QUAS-mCD8::GFP* transgenic mosquito stocks were generated as described in Riabinina et al. 2016 (6). *QUAS-GCaMP6f* was generated as described in Afify et al. 2019 (51). Wild-type Ngousso mosquitoes were a gift from the Insect Transformation Facility (Rockville, MD). *Flies. Orco-GAL4* (#26818) and *5xUAS-AgOr2* (#58828) lines were obtained from the Bloomington *Drosophila* Stock Center.

### Recombinant DNA construction

Plasmids were constructed by enzyme digestions, PCR, subcloning and the In-Fusion HD Cloning System (Clontech, catalogue number 639645). Plasmid inserts were verified by restriction enzyme digests and DNA sequencing. Insertions of each plasmid into the *Anopheles* genome (*QUAS-AgOr2, QUAS-mutAgOr2*) or the *Drosophila* genome (*20xUAS-AgOr2*) were verified by sequencing the vector-specific cassette within the transgenic animal.

To create the *pXL-BacII-15xQUAS-TATA-AgOr2-Sv40* reporter line, we linearized the *pXL-BacII-15xQUAS_TATA-Sv40* (6) vector with *Xhol*. The cDNA of *AgOr2* was amplified from Bloomington stock number 58828 using the oligos *Aga_OR2_F* (5’-ATTCGTTAACAGATCTATGCTGATCGAAGAGTGTCCGA-3’) and *Aga_OR2_R* (5’-CCTTCACAAAGATCGACGTCTTAGTTGTACACTCGGCGCAGC-3’). The resultant PCR product was then infusion-subcloned back into the construct. To create the *pXL-BacII-15xQUAS-TATA-mutAgOr2-Sv40* reporter line, we did a double digest of *pXL-BacII-15xQUAS-TATA-AgOr2-Sv40* with *BgIII* and *XhoI*. We then amplified *AgOr2* from *pXL-BacII-15xQUAS-TATA-AgOr2-Sv40* using a forward primer that mutated the start codon: ATG→TTT (*InfuMUTAgOr2_for*: 5’-ATTCGTTAACAGATCTTTTCTGATCGAAGAGTGTCCGATAATTG). We also engineered the reverse primer to create a frameshift mutation at a second in-frame ATG site located between the first and second transmembrane domains of *AgOr2* (*InfuMUTAGOr2_rev*: 5’-CGTCATTTTTCTCGAGTAGAGAGCGTACTCGGCGGC-3’).

The *20xUAS-AgOr2* reporter was created in *Drosophila* to test whether increasing the dosage of *AgOr2* affects olfactory physiology. The construct was made by digesting *pJFRC-20xUAS-IVS-CD8GFP* with *NotI* and *XbaI* and isolating the linearized 8.1kb vector. *AgOr2* was PCR amplified from *pXL-BacII-15xQUAS-TATA-AgOr2-Sv40* using the primers *UAS-AgOr2-FOR* (5’-TTACTTCAGGCGGCC GCAAA ATGCTGATCGAAGAGTGTCCG) and *UAS-AgOR2-REV* (5’-ACAAAGATCCTCTAGA TTAGTTGTACACTCGGCGCAG-3’). The PCR product was infusion cloned into the digested *pJRFRC-20xQUAS* vector. Upon sequence confirmation, the plasmid was midiprepped (Qiagen 12145) and sent to Rainbow Transgenics for injection into the attP site (RFT # 8622).

The *pXL-BACII-DsRed-OR7_9kbProm-QF2-hsp70* construct was used by the *Orco-QF2* driver line in this study. Construction of this plasmid is described in Riabinina et al. 2016 (6).

### *Anopheles gambiae* transgenics

*Anopheles gambiae* M-form strain Ngousso (the M-form of *An. gambiae* is now referred to as *Anopheles coluzzii*) mosquitoes were grown at 28°C, 70-75% relative humidity, 12h light/dark cycle. Freshly deposited eggs were collected by providing mated, gravid females with wet filter paper as an oviposition substrate for 15-20min, after which the eggs were collected and systematically arranged side-by-side on a double-sided tape fixed to a coverslip. Aligned embryos were covered with halocarbon oil (Sigma, series 27) and injected at their posterior pole with an injection cocktail between 30-40min after egg laying. Injection cocktails consisted of a mixture of two plasmids, one with a piggyBac vector carrying the transgene of interest with a dominant visible marker gene – enhanced cyan fluorescent protein (ECFP) – under the regulatory control of the *3xP3* promoter, and a piggyBac transposase-expressing plasmid consisting of the transposase open reading frame under the regulatory control of the promoter from the *An. stephensi vasa* gene. Vector concentrations were at 150ng/uL and the transposase-expressing plasmid was at 300ng/uL in 5mM KCL, 0.1mM sodium phosphate pH 6.8. Halocarbon oil was immediately removed and coverslips with injected embryos were placed in trays of water at 28°C, where the first instar larvae hatched ~24hr later. The Insect Transformation Facility (https://www.ibbr.umd.edu/facilities/itf) within the University of Maryland College Park’s Institute for Bioscience and Biotechnology Research performed all embryo microinjections. Adults developing from injected embryos were separated by sex at the pupal stage before mating, and small groups of 5-10 injected adult males or females were crossed to wild-type Ngousso adults of the opposite sex. The progeny from these matings were screened during the third or fourth larval instar for the presence of vector-specific marker gene expression. Transgenic larvae were saved and adults from these larvae were outcrossed to wild-type for a total of 5 generations.

### Insect stock maintenance

#### Anopheles gambiae

*Anopheles* mosquitoes were grown at 28°C, 70-75% relative humidity and 14hrlight/10hr dark cycle. Larvae were reared at low densities (175 larvae/1L dH_2_O) to ensure large adult size. They were provided with TetraMin Tropical Flakes and Purina Cat Chow Indoor pellets *ad libitum*. Pupae were hand collected and allowed to eclose in small cages, where they were provided with 10% sucrose continuously. Almost all pupae eclosed the day after collection. Adult males and females were kept together in the same cage for 7-10 days, after which they were fed mouse blood from anaesthetized mice according to Johns Hopkins University Animal Care and Use Committee (ACUC) approved protocol #M019M483. Eggs were collected from the resulting gravid females by providing them with a cup of water containing wet filter paper on which to deposit their eggs as an oviposition substrate. Each generation was screened for the presence of the eye specific marker encoded by the inserted plasmid cassette. *Drosophila melanogaster*. Flies were reared at 25°C and 70% humidity on a standard cornmeal diet.

### Calcium imaging

#### Preparation

*In vivo* preparation of mosquitoes (ages 3-10 days) and optical imaging of odor-evoked calcium responses are described in Afify et al. 2019 (51). *Genotyping mosquitoes*. After the recordings were made for each sample, we froze the bodies of all mosquitoes for subsequent gDNA extraction and genotyping. At the time of the experiment, our transgenic lines were not homozygous. Because all *QUAS* effector lines (*QUAS-AgOr2, QUAS-mutAgOr2, QUAS-GCaMP6f*) are marked with the dominant eye marker, ECFP, we had to determine – for each sample – whether the mosquito contained a single copy of *QUAS-AgOr2*, a single copy of *QUAS-GCaMP6f*, or both *QUAS-AgOr2* and *QUAS-GCaMP6f* transgenes (for experiments in **Fig. 2 and S1–2**). For the experiment in **Fig. 3**, we had to determine – for each sample – whether the mosquito contained a single copy of *QUAS-mutAgOr2*, a single copy of *QUAS-GCaMP6f*, or both *QUAS-mutAgOr2* and *QUAS-GCaMP6f* transgenes. To genotype *QUAS-GCaMP6f*, we used the primers *gcamp6f_for2* (5’-ATGGTATGGCTAGCATGACTG-3’) and *gcamp6f_rev* (5’-GTAGTTTACCTGACCATCCCC-3’). Females that did not have any amplification of *GCaMP6f* were discarded from the analysis. To genotype *QUAS-AgOr2* or *QUAS-mutAgOr2*, we used the following primers: *AgOr2_for1* (5’-TAATTGGTGTCAATGTGCGAG-3’) and *AgOr2_rev2* (5’-TTATCGGCTCCTCAAAGTCTG-3’). The PCR was designed so that both the wild-type *AgOr2* (1542bp) and the transgenic *AgOr2* (966bp) (or *mutAgOr2*; 968bp) – if present – would amplify. For each female, we determined whether she contained the wild-type and transgenic copy of *AgOr2* (or *mutAgOr2*) or only the wild-type *AgOr2*. Scoring of all calcium imaging files was done blind to genotype. *Analysis*. To make the heatmaps (ΔF), Fiji software (52) was used with a custom-built macro. This Macro uses the “Image stabilizer” plug-in to correct for movements in the recording, followed by the “Z project” function to calculate the mean baseline fluorescence (mean intensity in the first 9s of recording, before stimulus delivery). The “Image calculator” function was used to subtract the mean baseline fluorescence from the image of maximum fluorescence after odorant delivery (this image was manually chosen). Afterward, this ΔF image was used to produce heatmaps. To quantify the ΔF value for each segment to each odor, the “ROI manager” tool in Fiji was used to manually select an ROI. For each sample, we manually drew the ‘antennal ROI’ around the 11th antennal segment from an epifluorescent image taken for each sample prior to the calcium imaging. We also drew a ‘background ROI’ outside of the tissue using the same surface area as the ‘antennal ROI’ to control for any background signal. A “task manager” was used to store the location of the antennal ROI and the background, and the mean intensity across the antennal segment (or background control) for each odor was stored for each ROI. The final ΔF value was taken for each antennal segment and each odor as the mean intensity of the ‘antennal ROI’ minus the ‘background ROI.’ Of note, the imaging software was upgraded for **Fig. 3**, which explains why the (ΔF) values in **Fig. 3** are on a different scale than **Fig. 2, Fig S1**, and **Fig. S2**. Data were analyzed using JMP Version 9, SAS Institute Inc., Cary, NC, 1989-2019.

### Single Sensillum Recordings (SSR)

#### Drosophila preparation

Flies were housed on regular food in groups of a maximum of 10. Analysis was done on ab1 sensilla from 6-10 day old male flies. Sensilla of targeted ORNs were prepared and identified using methods described in Lin & Potter 2015 (53). Briefly, ab1 sensilla were identified by their response to CO_2_. Ab1 is the only sensillar group that houses the CO_2_-responsive neuron, and so a CO_2_-response was indicative that the sensillum we were recording from was ab1. Signals were amplified 100X (USB-IDAC System; Syntech, Hilversum, The Netherlands), inputted into a computer via a 16-bit analog-digital converter and analyzed off-line with AUTOSPIKE software (USB-IDAC System; Syntech). The low cutoff filter setting was 50Hz, and the high cutoff was 5kHz. To deliver odors or CO_2_ to ab1, a constant air stream was guided through a serological pipette with a tip placed 1cm from the antennae. The chemical cartridge was laterally inserted into this airflow. Stimuli consisted of 1000ms air pulses passed over odorant sources, which were various odorants diluted in paraffin oil (30μL on a filter paper of 1×2cm). For the benzaldehyde analysis, we counted every spike (did not distinguish among neuron subtypes). For the methyl salicylate analysis, we only included the response from the DmOR10a-positive neuron. Delta spikes/second were calculated by manually counting the number of spikes in a 0.5s window at stimulus delivery (200ms after stimulus onset to account for the delay due to the air path) and then subtracting the number of spontaneous spikes in a 0.5s window before stimulation, multiplied by 2 to obtain delta spikes per second. *Mosquito preparation*. Mated females were 5-12 days old and not bloodfed. Extracellular recordings of the capitate pegs on the maxillary palps were made using the same equipment as for *Drosophila* SSR (see above). Cp sensilla were also identified by their response to CO_2_. Cp is the only sensillum that houses the CO_2_-responsive neuron, and so a CO_2_-response was indicative that the sensillum we were recording from was cp. SSR data were analyzed using JMP Version 9, SAS Institute Inc., Cary, NC, 1989-2019.

### RNA-seq

To generate experimental pools of *Orco>AgOr2* antennae, mosquitoes of the genotype *QUAS-AgOr2* were crossed to *Orco-QF2*. Simultaneously, we generated control samples by crossing our *Orco>AgOr2* or *Orco-QF2* strains to wild-type (2 crosses). All 3 crosses were conducted in large breeding cages that consisted of ~75 males and ~75 females. Crosses were blood-fed a total of 4 times and progeny from each cross were screened for the eye-specific fluorescent markers. Larvae generated from experimental crosses were screened for the presence of ECFP and DsRed, markers for the *QUAS* and the *Orco-QF2* transgenes, respectively. Larvae that did not contain both markers were discarded. For the control larvae progeny, animals that contained a single eye marker (either DsRed or ECFP, respectively) were kept in control pools that consisted either of *Orco-QF2* or *QUAS-AgOr2* alone. All mosquitoes used in this study were heterozygous for a given transgene. *Orco>mutAgOr2* and *QUAS-mutAgOr2* library preparations were made using the procedure described above with the exception that the antennae were extracted at a different time. To prepare antennal RNA-seq libraries, we isolated RNA from approximately 200 antennae from age-matched cohorts (11 total samples: 3 experimental samples containing *Orco>AgOr2* antennae (*orco-QF2* + *QUAS-AgOr2*), 5 control samples consisting of 2 samples from *Orco-QF2* antennae, 2 samples from *QUAS-AgOr2* antennae, and 1 sample from *QUAS-mutAgOr2* antennae, and 3 experimental samples containing *Orco>mutAgOr2* antennae). These mated females were within their fertile period (5-20 days old) and did not receive a bloodmeal.

To create the antennal RNA-seq libraries, we removed the whole antennae from the base of the pedicel and isolated total RNA using TriZol purification methods. The tissues were disrupted and homogenized using a power pestle with disposable RNAse free pestles. Total RNA samples were stored at −80°C and shipped to Genewiz Inc, where they were first assessed for quantity (Qubit Quantification) and quality (Agilent 2100 Bioanalyzer). RNA library preparation with polyA selection was then carried out using the *Illumina* HiSeq with a 2×150bp configuration. Paired end reads were pseudoaligned to the AgamP4.12 reference transcriptome (7) and isoform-level abundances were quantified using kallisto v0.46.0 (54) using default parameters with the following exceptions: `-t4 and -b100`. Per-sample abundances were aggregated and normalized in R using sleuth v0.30.0 (55). For each pairwise comparison (AgOr2 v control, mutAgOr2 v control, and mutAgOr2 v AgOr2), we fit a full sleuth model using condition as an explanatory variable. Differentially expressed isoforms were identified using the Wald test (q≤0.01, Benjamini-Hochberg corrected) for each model fit. Ternary plots were constructed using the ggtern R package (56).

The RNA-seq raw reads and dataset analyses are available from NCBI. The accession numbers are: (TBD)

### Immunohistochemistry

#### Brains

Brains of female mosquitoes of *Orco>mCD8::GFP* and *Orco>AgOr2,mCD8::GFP* genotypes were extracted and stained as described previously (6). Once genotyped via a PCR designed to amplify the transgenic and wild-type copies of *AgOr2* (see ‘Calcium imaging section’ above), experimental and control brains were separated and stained in two different groups. Rat anti-CD8 (Invitrogen #MCD0800, 1:100) was used to visualize the ORN projections to the antennal lobes and mouse nc82 (DSHB, 1:50) was added to visualize the structure of the brain. *Pupal Antennae. Orco>mCD8::GFP* larvae were collected in 2 trays, each of which contained ~60 larvae. At the given timepoint After Puparium Formation (APF), the cephalothorax and abdomen of the pupae were extracted and the antennae, oculus, compound eye, and rudimentary appendages were placed in 4% paraformaldehyde (PFA) with 0.1% Triton Phosphate Buffer Solution (PBS-T). Antennae were then dissected out and washed 3x in 1X PBS-T (0.1%). Antennae were then blocked for 30min at 4°C with 5% Normal Goat Serum (NGS).

To visualize mCD8::GFP expression, we added rat anti-CD8 (Invitrogen #MCD0800, 1:100), which was left to incubate on a rotator overnight at 4°C. The next day, antennae were washed 3x in 1X PBS-T (0.1%) and Alexa-488 goat anti-rat (Invitrogen A110066, 1:200) was added to the solution. After 1.5hr at 25°C, antennae were washed 2X in 1X PBS-T (0.1%) and the last wash in 1xPBS to remove the triton. Antennae were placed on slides and mounted in Slow Fade Gold medium (ThermoFisher Scientific #S36936).

### Confocal imaging and analyses

Brains and pupal antennae were imaged on an LSM 700 Zeiss confocal microscope at 512×512 pixel resolution, with 0.96 or 2.37uM Z-steps. For illustration purposes, confocal images were processed using Fiji (52) to collapse the maximum intensity projection of Z-stacks into a single image. We used cells that were immunoreactive to mCD8 as a proxy for ORCO expression. Cells were counted manually.

### Host-proximity assay

The design and methods of the host-proximity assay are modified from DeGennaro et al. 2013 (4). Briefly, for each trial, 20-30 adult female *Anopheles* mosquitoes (5+ days post eclosion, mated but not blood-fed) were sorted under cold anesthesia (4°C), placed in 24-oz deli containers (https://www.amazon.com/gp/product/B00NB9WCEO/ref=oh_aui_detailpage_o06_s00?ie=UTF8&psc=1), and fasted with access to water for 16-24hr prior to assaying. Pre-fasting and behavior experiments took place at 27°C and 70-80% relative humidity. Experiments took placed after Zeitgeber Time 0 (ZT0) and continued until ZT8. 5 minutes before the start of the assay, mosquitoes were released into a modified BugDorm (https://shop.bugdorm.com/bugdorm-1-insect-rearing-cage-p-1.html). After 5 minutes had elapsed, pure CO_2_ and synthetic air (air flow rate: 0.1L/min, CO_2_ flowrate 145MM) were mixed in an adaptor before being pulsed (3sec on: 3 sec off) into a flypad (8.1 x 11.6cml catalogue #59-114; Flystuff.com, San Diego, CA), which was placed at the bottom of the BugDorm. Mosquitoes were then presented with a single volunteer’s arm, which was placed 2.5cm away from one side of the BugDorm so that mosquitoes could not come into direct contact with the arm. To control the distance from the arm to the cage, a Q-Snap needlework frame (https://www.amazon.com/dp/B00013MV30/ref=twister_B07CQQJKL2?_encoding=UTF8&psc=1) was placed flush against the cage and the arm was pressed against the vials. The arm was elevated 2.7cm by placing it on a plastic microcentrifuge test-tube rack. An HDR-CX260V camera (Sony) was positioned to take images of mosquitoes responding to the human arm. Trials ran for 3 minutes. To quantify mosquito responses that came into ‘close proximity’ to the human arm, we counted the number of mosquitoes resting on the screen. We did not include mosquitoes that had landed on the white area surrounding the screen nor the mosquitoes that were in flight. For the % attraction figure, we scored the number of mosquitoes that came into close proximity of the arm every 10 seconds from minute 1 to minute 3 and then divided this number by the total number of mosquitoes in the trial. Data were analyzed using JMP Version 9, SAS Institute Inc., Cary, NC, 1989-2019.

## Author Contributions

S.E.M and C.J.P. designed research; S.E.M., A.A., and L.A.G. performed research; S.E.M. and L.A.G. analyzed data; and S.E.M. and C.J.P. wrote the paper.

## Acknowledgments

This research benefited from the limitless enthusiasm and support of Potter Lab members, both past and present. In particular, we would like to thank Kateline Robinson Shaw and Liz Marr for assisting with molecular biology. Darya Task and Joanna Konopka helped rear the transgenic mosquitoes and improved the writing of the manuscript. We also thank Greg Artiushin for the graphic designs he made for the paper. This work was supported by grants from the National Institutes of Health to C.J.P. (NIAID R01Al137078), the Department of Defense to C.J.P. (W81XWH-17-PRMRP), and a Johns Hopkins Malaria Research Institute Postdoctoral Fellowship to SEM. We thank the Johns Hopkins Malaria Research Institute and Bloomberg Philanthropies for their support.

---

**Supplementary Figure 1.**
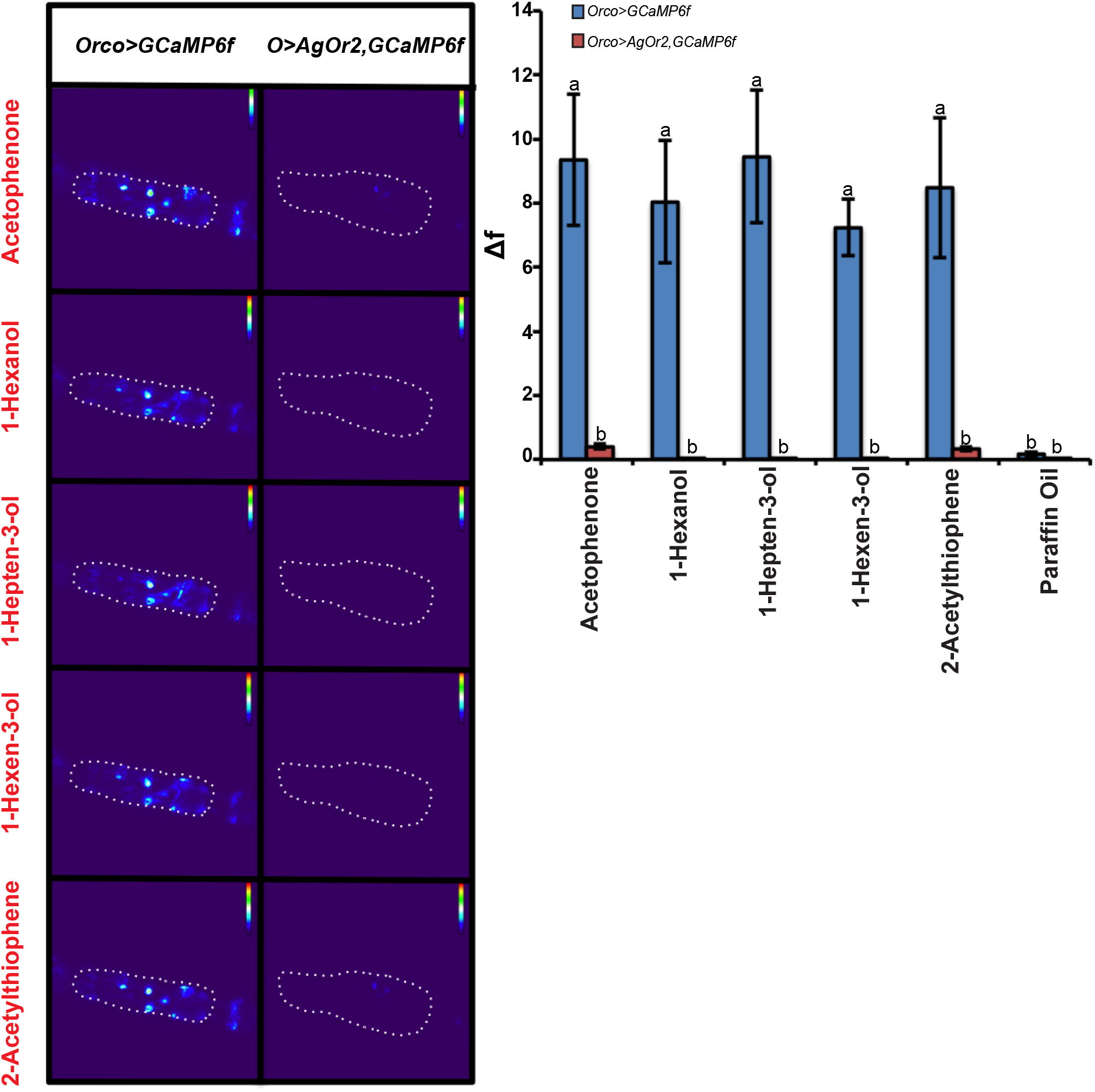
Antennae with ectopic *AgOr2* expression fail to respond to odors that activate multiple ORN classes. The activity of olfactory receptor neurons in antennal segment 11 (outlined by white dots) towards the listed odors were detected by calcium imaging of *Orco*-positive neurons expressing GCaMP6f (51). *Orco>AgOr2,GCaMP6f* is listed here as *O>AgOr2,GCaMP6f*. All odors were presented at a 10% concentration. A two-way repeated measures ANOVA was used to determine that there was a significant effect at the *p*<0.005 level of odor and genotype on calcium responses. Groups with different letter values (a-b) are statistically different as determined by the Tukey post hoc HSD test. According to Carey et al. 2010 (16), acetophenone, 1-hexanol, 1-hepten-3-ol, 1-hexen-3-ol, 2-acetylthiophene activate 16, 14, 16, 14, 15 of the tested ORs, respectively, at a rate of >50 spikes/sec. Each sample included in the analysis was taken from a different female mosquito. n_*Orco>GCaMP6f*_ = 5, n_*Orco>AgOr2,GCaMP6f*_ = 9.

**Supplementary Figure 2.**
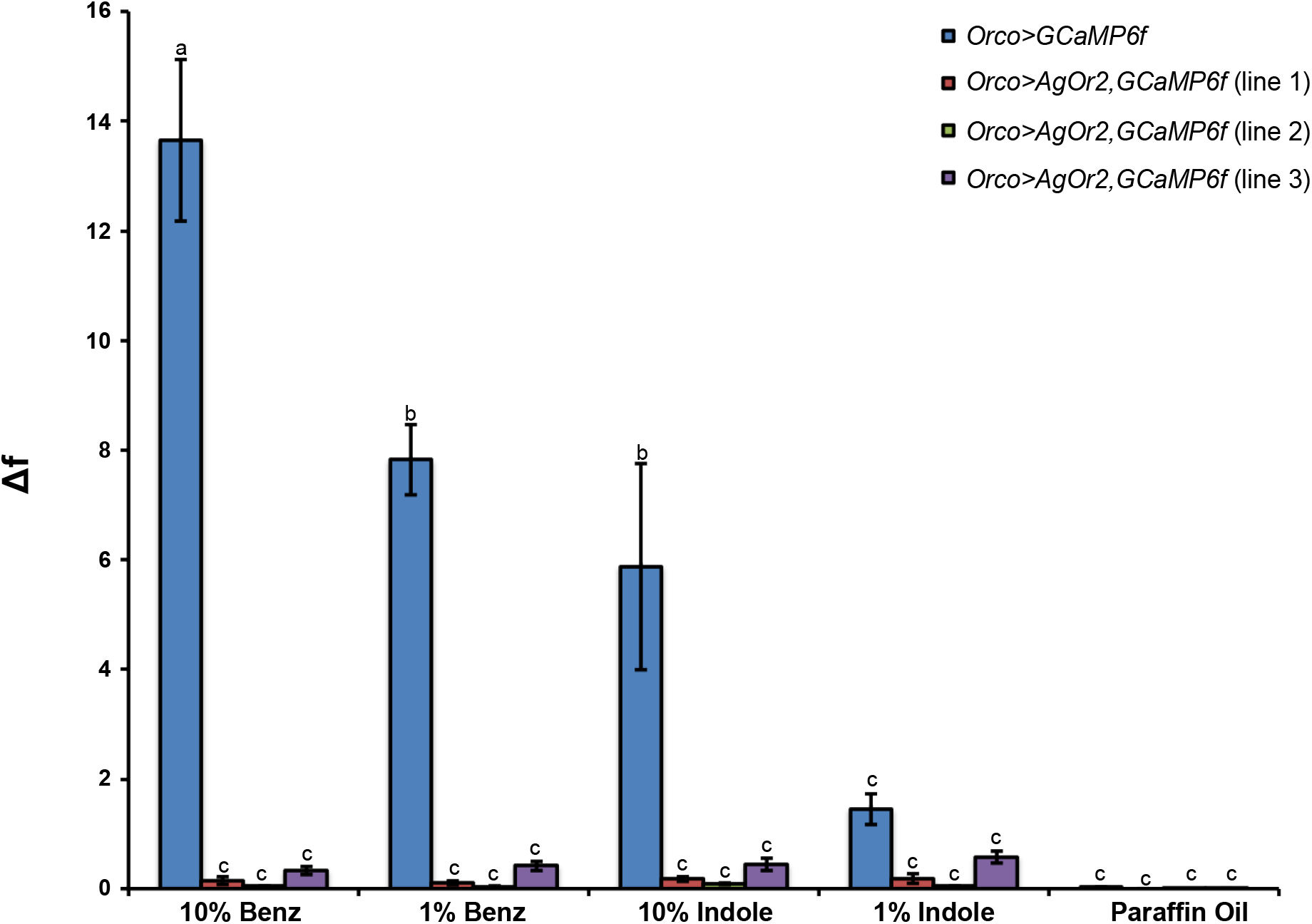
The dominant negative phenotype of *Orco>AgOr2* is independent of the *QUAS-AgOr2* insertion site into the genome. Two additional *QUAS-AgOr2* lines (lines 2 and 3) show impaired physiology in the presence of benzaldehyde and indole when compared to wild-type. A two-way repeated measures ANOVA was used to determine that there was a significant effect of genotype and odor on calcium responses at the *p<0.0001* level. Groups with different letter values (a-c) are statistically different as determined by the Tukey post hoc HSD test. Each sample included in the analysis was taken from a different female mosquito. n_*Orco>GCaMP6f*_ = 9; n_*Orco>AgOr2,GCaMP6f (line 1)*_ = 8; n_*Orco>AgOr2,GCaMP6f (line 2)*_ = 7; n_*Orco>AgOr2,GCaMP6f (line 3)*_ = 9.

**Supplementary Figure 3.**
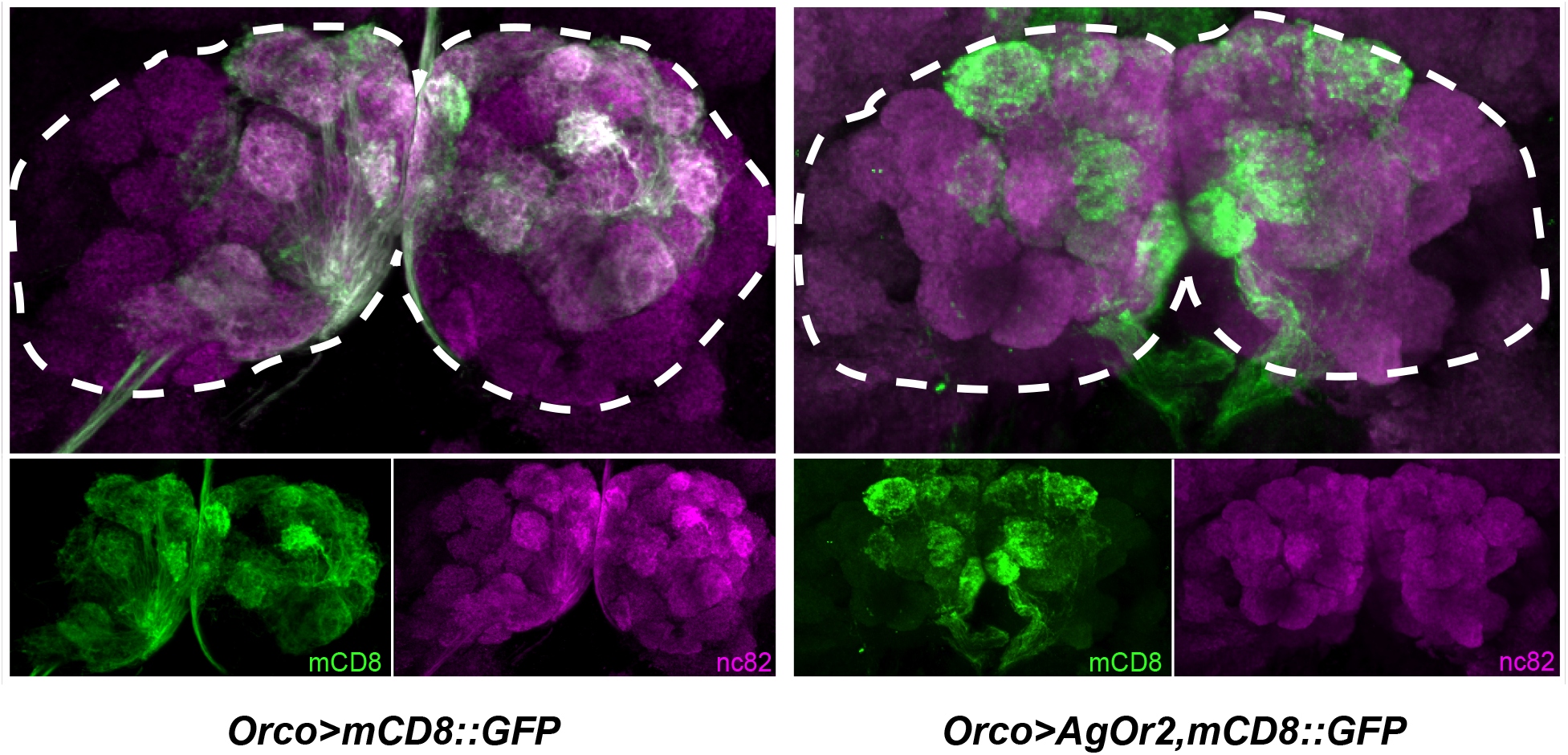
*Orco*-positive neuron processes are present in the adult *Orco>AgOr2,mCD8::GFP* antennal lobe. *Orco*-positive ORNs send their projections to the antennal lobe of control (*Orco>mCD8::GFP*) and experimental (*Orco>AgOr2, mCD8::GFP*) lines. Anti-nc82 was used to visualize the structure of the antennal lobes and anti-CD8 was used to visualize ORN projections. Antennal lobes are outlined with white dotted lines. n_*Orco>mCD8::GFP*_ = 9; n_*Orco>AgOr2,mCD8::GFP*_ = 2.

**Supplementary Figure 4.**
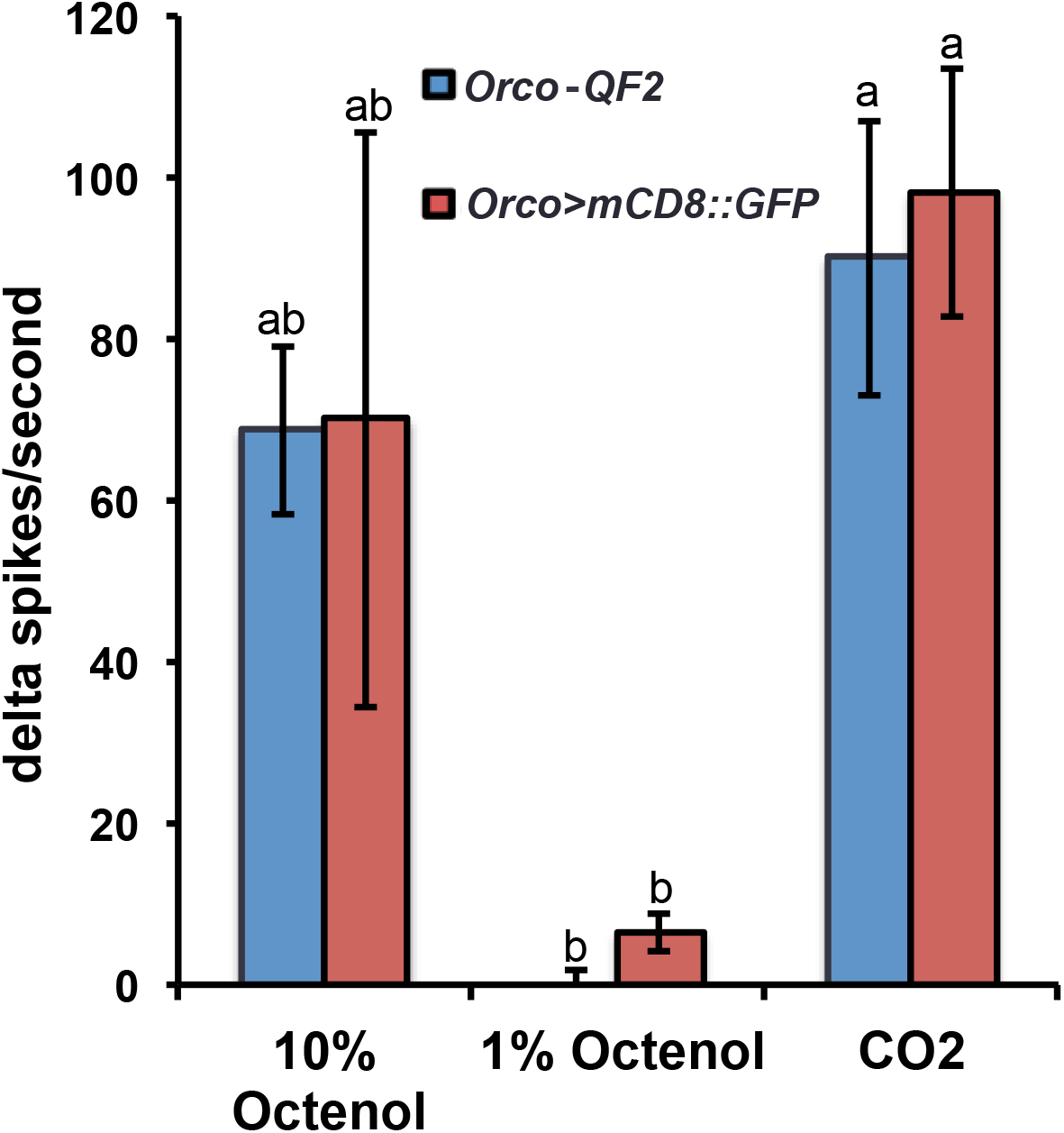
Driving a generic membrane-bound protein (*mCD8::GFP*) does not impair *Orco*-positive responses to odor. Single sensillum recordings were performed in the maxillary palp capitate peg sensilla. There was no difference in how cpB/C neurons of *Orco>mCD8::GFP* and *Orco-QF2* genotypes responded to 10% and 1% octenol. The presence of the *Orco*-negative CO_2_ neuron was used to verify that we were recording from a cp sensillum. A two-way repeated measures ANOVA was used to determine that there was a significant effect at the *p*<0.005 level of odor but not genotype on neuronal responses. Groups with different letter values (a-b) are statistically different as determined by the Tukey post hoc HSD test. 2-3 Females *per* genotype were analyzed. The number of sensilla evaluated for each group: n*_Orco-QF2_* = 6; n_*Orco>mCD8::GFP*_ = 5.

**Supplementary Figure 5.**
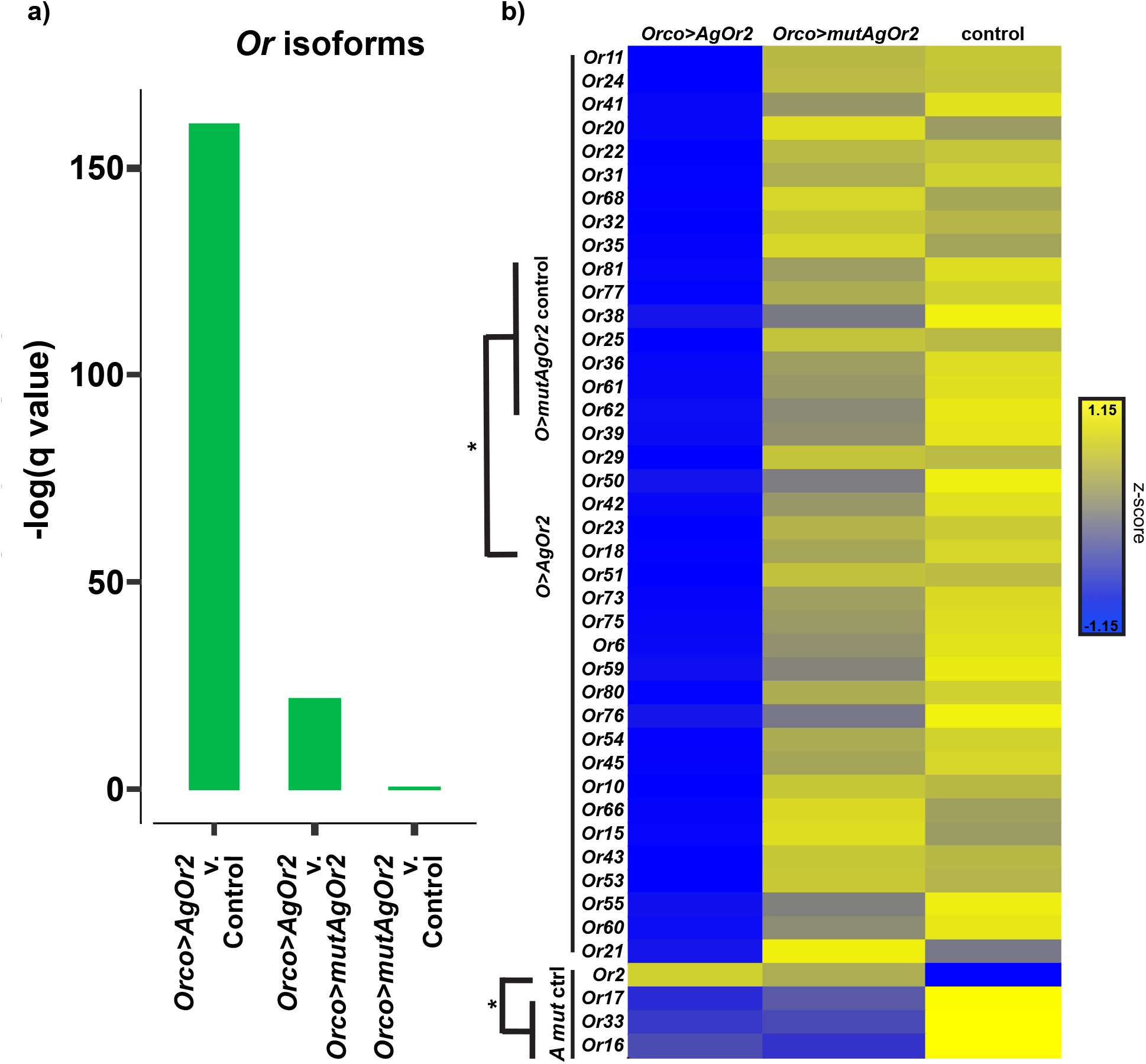
The majority of *Ors* are downregulated in *Orco>AgOr2* but not *Orco>mutAgOr2* and control genotypes. Reads from 3 *Orco>AgOr2*, 3 *Orco>mutAgOr2*, and 5 control samples were aligned to the *Anopheles gambiae* geneset AgamP4.12 (7) and samples were averaged for each group. (**a**) Results from the gene set enrichment test show significant differences in *Or* isoform levels between *Orco>AgOr2* v. control and *Orco>AgOr2 v. Orco>mutAgOr2*. (**b**) A heatmap was used to visualize the results of pair-wise tests comparing the average isoform abundance levels among *Orco>AgOr2, Orco>mutAgOr2*, and control groups. The first 39 *Ors* listed change between control and *Orco>AgOr2* but not control and *Orco>mutAgOr2*. The last 4 *Ors (Or2, Or17, Or33*, Or16) are the same between *Orco>AgOr2* (listed as ‘*A*’) and *Orco>mutAgOr2* (listed as ‘*mut*’) but different in control (listed as ‘ctrl’). *Ors* not included in the heatmap show no differences among groups. For each cell, a z-score was calculated by subtracting the mean isoform TPM from the cell’s TPM divided by the standard deviation of the isoform TPM.

**Supplementary Figure 6.**
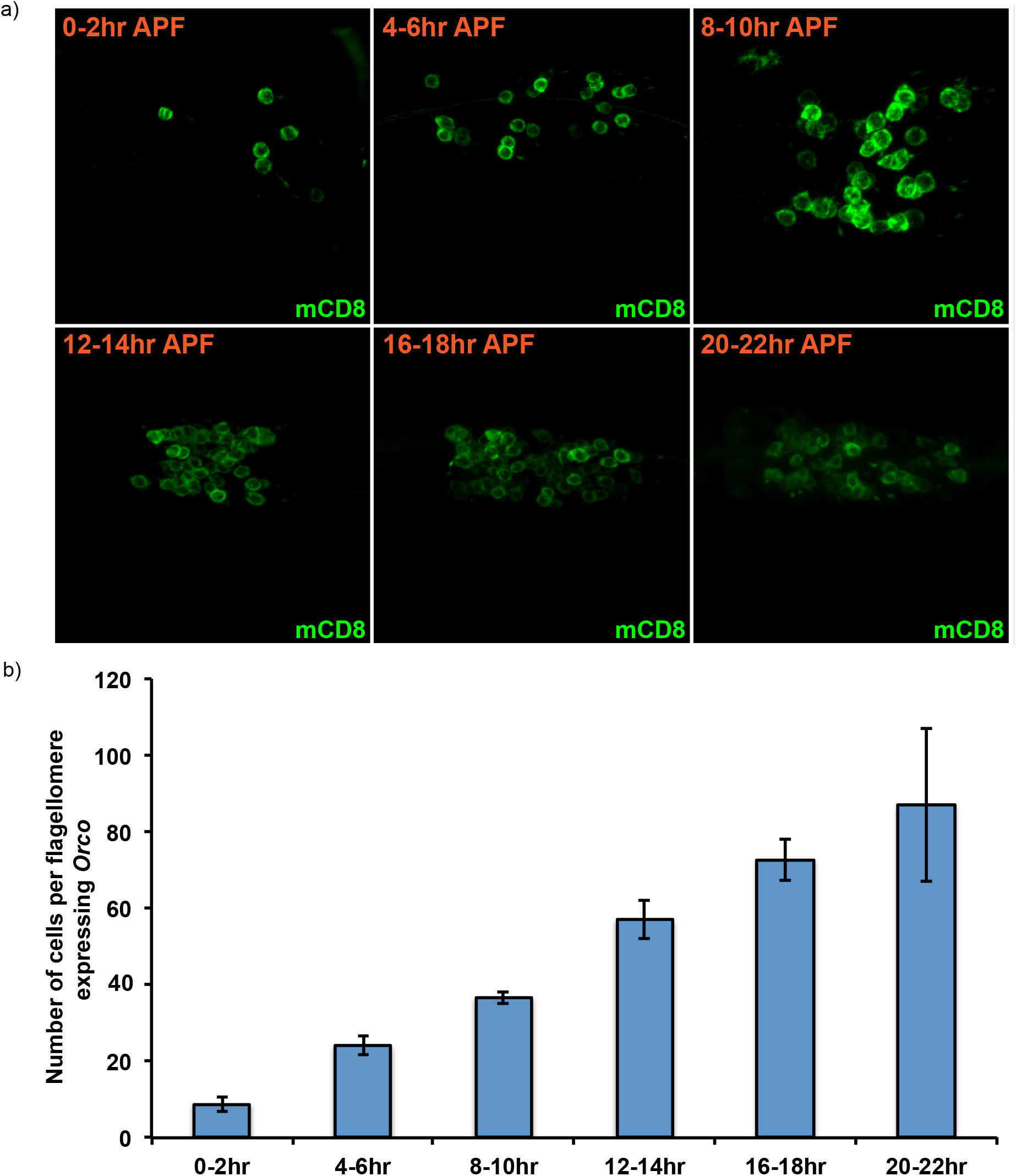
ORCO is expressed at the start of pupal ecdysis. **a)** Representative images of ORCO-expressing neurons in the *Orco>mCD8::GFP* genotype. Pupal antennae were extracted at the given timepoint After Puparium Formation (APF) and stained for anti-mCD8 (green). **b)** Cells were scored as ORCO-positive based on the presence of mCD8. Cells from one flagellomere per animal in an average of 5 animals *per* timepoint were scored. The flagellomere that was scored was randomized for each sample. The error bars represent standard deviation of the mean.

